# Community-curated and standardised metadata of published ancient metagenomic samples with AncientMetagenomeDir

**DOI:** 10.1101/2020.09.02.279570

**Authors:** James A. Fellows Yates, Aida Andrades Valtueña, Ashild J. Vågene, Becky Cribdon, Irina M. Velsko, Maxime Borry, Miriam J. Bravo-López, Antonio Fernandez-Guerra, Eleanor J. Green, Shreya L. Ramachandran, Peter D. Heintzman, Maria A. Spyrou, Alexander Hübner, Abigail S. Gancz, Jessica Hider, Aurora F. Allshouse, Christina Warinner

## Abstract

Ancient DNA and RNA are valuable data sources for a wide range of disciplines. Within the field of ancient metagenomics, the number of published genetic datasets has risen dramatically in recent years, and tracking this data for reuse is particularly important for large-scale ecological and evolutionary studies of individual microbial taxa, microbial communities, and metagenomic assemblages. AncientMetagenomeDir (archived at https://doi.org/10.5281/zenodo.3980833) is a collection of indices of published genetic data deriving from ancient microbial samples that provides basic, standardised metadata and accession numbers to allow rapid data retrieval from online repositories. These collections are community-curated and span multiple sub-disciplines in order to ensure adequate breadth and consensus in metadata definitions, as well as longevity of the database. Internal guidelines and automated checks to facilitate compatibility with established sequence-read archives and term-ontologies ensure consistency and interoperability for future meta-analyses. This collection will also assist in standardising metadata reporting for future ancient metagenomic studies.

## Background & Summary

A crucial but often overlooked component of scientific reproducibility is the efficient retrieval of sample (meta)data. While the field of ancient DNA (aDNA) has been celebrated for its commitment to making sequencing data available through public archives^1^, the retrieval of this data is not always trivial. The field of ancient metagenomics benefits from large sample sizes and the ability to reuse previously published datasets. However, the current absence of standards in basic metadata reporting can make data retrieval tedious and laborious, leading to analysis bottlenecks.

Ancient metagenomics can be broadly defined as the study of the *total* genetic content of temporally-degraded samples^2^. Areas of study that fall under ancient metagenomics include studies of host-associated microbial communities (e.g., ancient microbiome studies of dental calculus or paleofeces^3^), genome reconstruction and analysis of specific microbial taxa (e.g., ancient pathogens^4^), and environmental reconstructions using sedimentary ancient DNA (sedaDNA)^5^. Genetic material obtained from ancient samples has undergone a variety of degradation processes that can cause the original genetic signal to be overwhelmed by modern contamination, requiring large DNA sequencing efforts to detect, quantify, and authenticate the remaining truly aDNA^6,7^. These studies have only become feasible since the development of next-generation sequencing (NGS), which employs massively parallel sequencing to generate large amounts of data that are mostly uploaded to and stored on large generalised archives such as the European Bioinformatic Institute’s (EBI) European Nucleotide Archive (ENA, https://www.ebi.ac.uk/ena/) or the US National Center for Biotechnology Information (NCBI)’s Sequence Read Archive (SRA, https://www.ncbi.nlm.nih.gov/sra). However, because these are generalised databases used for many kinds of genetic studies, searching for and identifying ancient metagenomic samples can be difficult and time consuming, partly because of the absence of standardised metadata reporting for ancient metagenomics data. Consequently, researchers must resort to repeated extensive literature searches of heterogeneously reported and inconsistently formatted publications to locate ancient metagenomics datasets. Overcoming the difficulty of finding previously published samples is particularly pertinent in studies of aDNA, as palaeontological and archaeological samples are by their nature limited and avoiding repeated or redundant sampling is a high priority^8^.

To address these issues, we established AncientMetagenomeDir, a community-curated collection of annotated sample lists that aims to guide researchers to all published ancient metagenomics-related samples. AncientMetagenomeDir was conceived by members of a recently established international and open community of researchers working in ancient metagenomics (Standards, Precautions and Advances in Ancient Metagenomics, or ‘SPAAM’ - spaam-workshop.github.io), whose aim is to foster research collaboration and define standards in analysis and reporting. The collection aims to be comprehensive but lightweight, consisting of tab-separated value (TSV) tables for three major sub-disciplines of ancient metagenomics. These tables contain essential, sample-specific information in the field of aDNA studies, including: geographic coordinates, temporal data, sub-discipline-specific critical information, and accession codes of public archives that guides researchers to associated sequencing data (see Methods). Keeping these tables in a simple format, and together with our comprehensive contribution guides, encourages continuous contributions from the community and facilitates usage of the resource by researchers coming from non-computational backgrounds, something common in interdisciplinary fields such as archaeo- and palaeogenetics.

AncientMetagenomeDir is designed to track the development of ancient metagenomics through regular releases. As of release v20.09, this includes 87 publications since 2011, representing 443 ancient host-associated metagenome samples, 269 ancient microbial genomes, and 312 sediment samples (Fig. 1), spanning 49 countries (Fig. 2). We expect Ancient-MetagenomeDir to deliver three key benefits. First, it will contribute to the longevity of important cultural heritage by guiding future sampling-strategies; reducing the risk of repeated or over-sampling of the same samples or regions. Second, it can form a starting point for the development of software to allow rapid aggregation and field-specific data processing. Finally, as a community-curated resource designed specifically for widespread participation, AncientMetagenomeDir will help the field to define a common standard of metadata reporting (such as with MIXs checklists^9^), facilitating the creation of further rich but consistent sample databases for future researchers.

**Figure 1.**
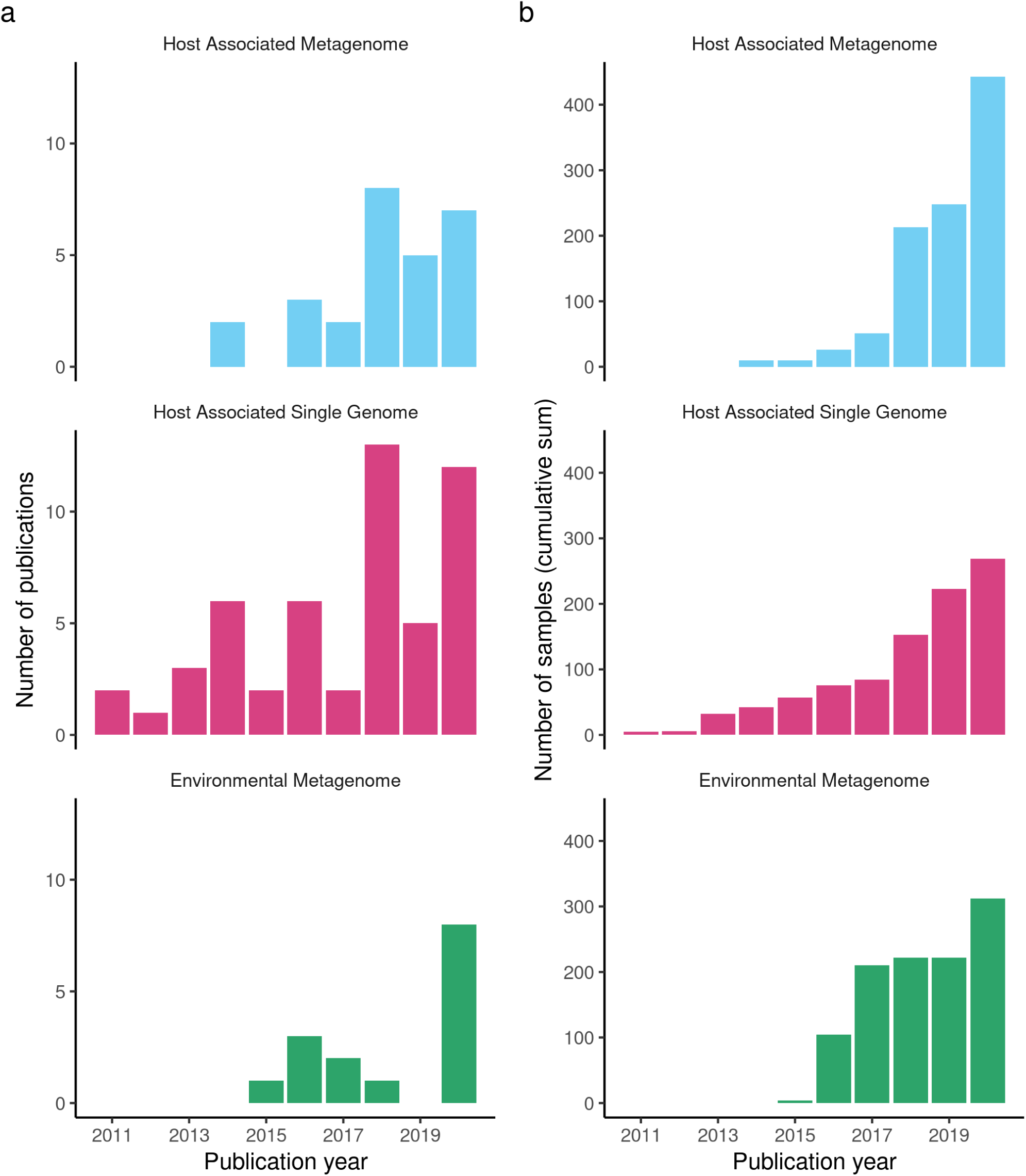
Timelines depicting the development of sub-fields of ancient metagenomics as recorded in AncientMetagenomedir as per release v20.09. (**a**) Number of ancient metagenomic publications per year. (**b**) Cumulative sum of published samples with genetic sequencing data or sequences in publicly accessible archives.

**Figure 2.**
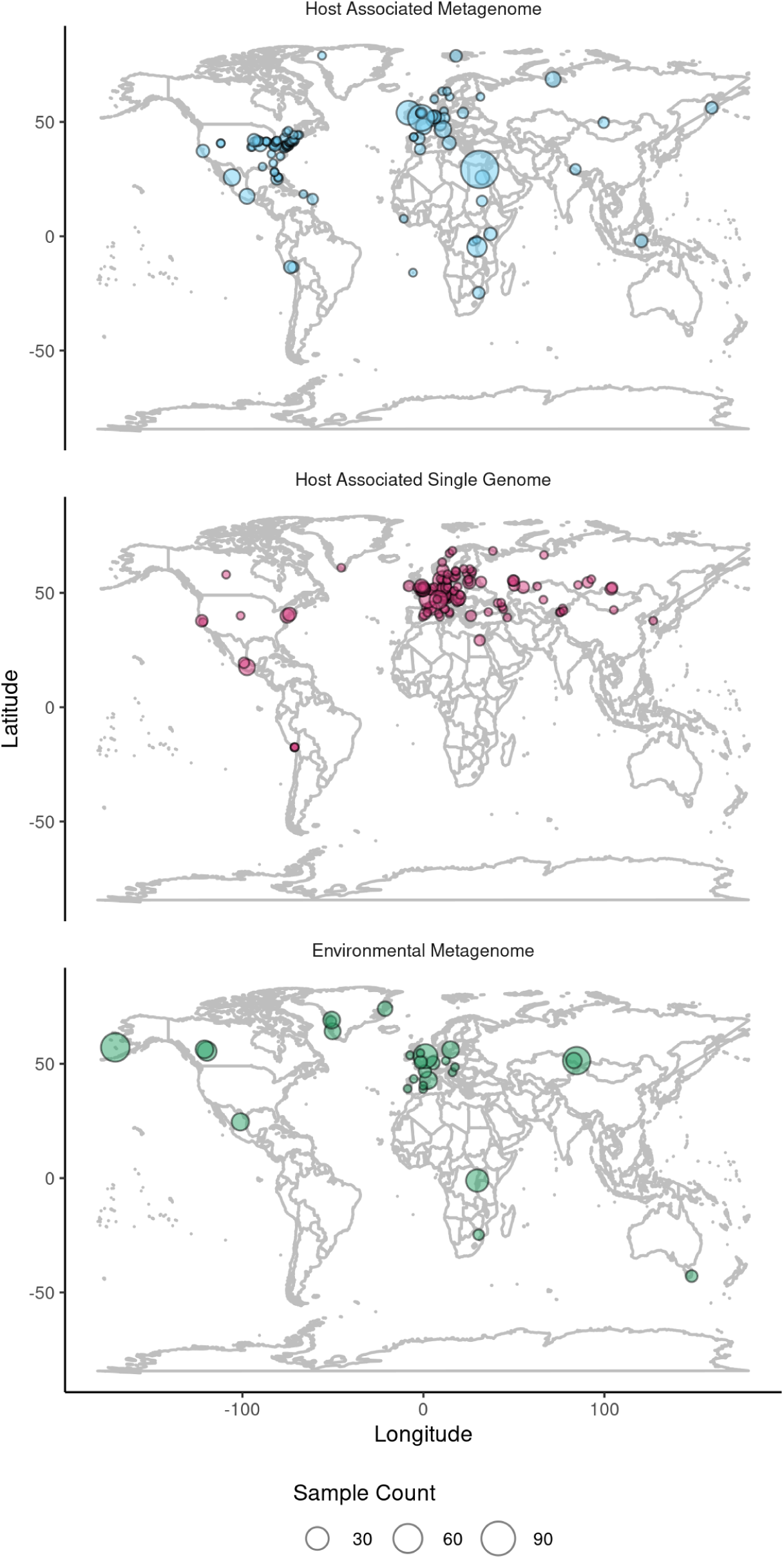
Maps depicting the geographic spread of samples with latitude and longitude information, between sub-fields of ancient metagenomics as recorded in AncientMetagenomeDir as per release v20.09.

## Methods

### Repository Structure

AncientMetagenomeDir^10^ is a community-curated set of tables maintained on GitHub, that contains metadata from published ancient metagenomic studies (https://github.com/SPAAM-workshop/AncientMetagenomeDir). While most submissions are made by SPAAM members, anyone with a GitHub account is welcome to propose and/or add publications for inclusion. Submitted publications must be published in a peer-reviewed journal; the purpose of AncientMetagenomeDir is not to act as a quality filter and it currently does not make assessments based on data quality. The tables are formatted as tab-separated value (TSV) files in order to maximize accessibility for all researchers and to allow portability between different data analysis software. Valid samples for inclusion currently fall under three categories: (1) host-associated metagenomes (i.e., host-associated or skeletal material microbiomes), (2) host-associated single genomes (i.e., pathogen or commensal microbial genomes), and (3) environmental metagenomes (e.g., sedaDNA). In addition, a fourth category is currently planned: (4) anthropogenic metagenomes (e.g., dietary and microbial DNA within pottery crusts, or microbial DNA and handling debris on parchment). To be included, samples must have been sequenced using a shotgun metagenomic or genome-level enrichment approach, and sequence data must be publicly available on an established or stable archive. Publications included in the current release were selected for inclusion based on direct contributions by authors and literature reviews. Publications are initially added as a GitHub ‘Issue’. Publications may belong to multiple categories, and the corresponding issue is tagged with relevant category ‘labels’ to assist with faster evaluation and task distribution.

### Data Acquisition

After an Issue (i.e., publication) is suggested, any member of the open SPAAM community can assign themselves to the Issue. The member then creates a git branch off of the main repository, manually extracts the relevant metadata from the given publication, and adds it to the corresponding table (e.g. host associated metagenome, or environmental metagenome). Extensive documentation is available to assist contributors to ensure correct entry of metadata, with one README file per table that contains column definitions and guidelines on how to interpret and record metadata. Extensive documentation on submissions, including instructions on using GitHub, are available via tutorial documents and the associated repository wiki.

The metadata in each table covers five main categories: publication metadata (project key, year, and publication DOI), geographic metadata (site name, coordinates, and country), sample metadata (sample name, material type, and (meta)genome type) and sequencing archive information (archive, sample archive accession ID). Due to inconsistency in the ways metadata are reported in publications and archives, and to maintain concise records we have specified (standardised) approximations for the reporting of sample ages, geographic locations, and archive accessions, following where possible MIxS^9^ categories. This approach allows researchers to access sufficiently approximate information during search queries to identify samples of interest (e.g., 3700 BP), which they can subsequently manually check to obtain the exact specifications reported in the original publication (e.g., Late Bronze Age, 3725 +/- 15 BP). Geographic coordinates are restricted to a maximum of three decimals, with fewer decimals indicating location uncertainty (e.g., if a publication only reports a province rather than a specific site). Dates are reported (where possible) as uncalibrated years Before Present (BP, i.e., from 1950), and rounded to the nearest 100 years, due to the range of calculation and reporting methods (radiocarbon dating vs. historical records, calibrated vs. uncalibrated radiocarbon dates, etc.). For sequence accession codes, we opted for using *sample* accession codes rather than direct sequencing data IDs. This is due to the myriad of ways in which data are generated and uploaded to repositories (e.g., one sample accession per sample vs. one sample accession per library; or uploading raw sequencing reads vs. only consensus sequences). We found that in most cases sample accession codes are the most straightforward starting points for data retrieval. However, we did observe errors in some data accessions uploaded to public repositories, such as multiple sample codes assigned to different libraries of the same sample, and insufficient metadata to link accessions to specific samples reported in a study. Overall, we found that heterogeneity in sample (meta)data uploading was a common problem, which highlights the need for improvements in both training and community-agreed standards for data sharing and metadata reporting in public repositories. In addition to metadata recorded across all sample types, we have added table-specific metadata fields to individual categories as required (e.g., species for single genomes and community type for microbiomes). Such fields can be further extended or modified with the agreement of the community.

### Data Validation

After all metadata has been added, a contributor makes a Pull Request (PR) into the master branch. Every PR undergoes an automated continuous-integration check via the open-source companion tool AncientMetagenomeDirCheck^11^ (https://github.com/SPAAM-workshop/AncientMetagenomeDirCheck, License: GNU GPLv3). In order to ensure consistency within and between metadata fields, this tool checks that the entries for each column match a given regex or category defined in controlled JSON ‘enum’ lists (stored in an ‘assets’ directory in the repository). For example, valid country codes are guided by the International Nucleotide Sequence Database Collaboration (INSDC) controlled vocabulary (http://www.insdc.org/country.html), host and microbial species names are defined by the NCBI’s Taxonomy database (https://www.ncbi.nlm.nih.gov/taxonomy), and material types are defined by the ontologies listed on the EBI’s Ontology Look Up service (https://www.ebi.ac.uk/ols/index) - particularly the Uberon^12^ and Envo ontologies^13,14^. These controlled vocabularies, alongside stable linking (via DOIs), ensures reliable querying of the dataset, and allows future expansion to include richer metadata by linking to other databases. Descriptions for the minimum required fields for an AncientMetagenomeDir table are provided in Table 1.

**Table 1.**
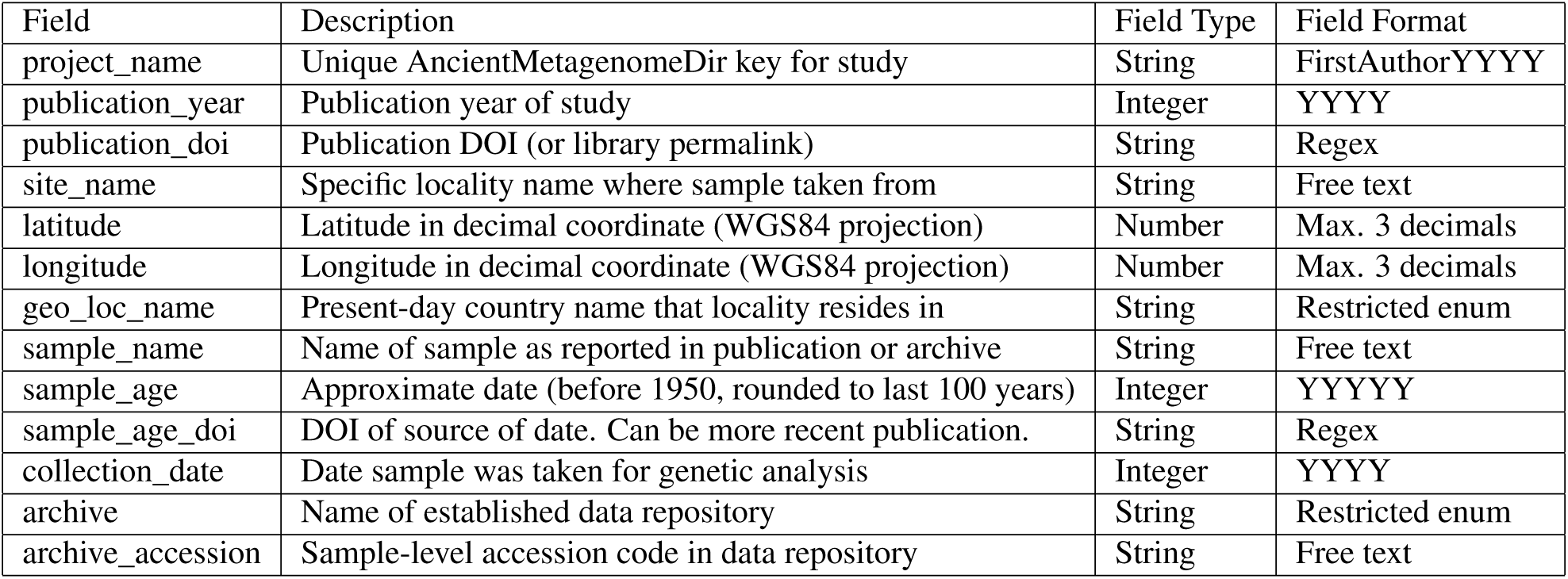
Core fields that are required for all AncientMetagenomeDir sub-discipline tables, including field type and standardised formatting description. Field formats are defined in a JSON schema, against which each new study submission is cross-checked. Further sub-discipline specific fields are included in the corresponding table, as required by the community.

Once automated checks are cleared, a contributor then requests a minimum of one peer-review performed by another member of the SPAAM community. This review involves checking the entered data for consistency against the table’s README file and also for accuracy against the original publication. Once automated checks and the peer-review are both passed, the publication’s metadata are then added to the master branch and the corresponding Issue is closed. For each added publication, a CHANGELOG is maintained to track the papers included in each release and to record any corrections that may have been made (e.g., if new radiocarbon dates are published for previously entered samples). The CHANGELOG or Issues pages on GitHub can be consulted to check whether a given publication has already been added (or excluded) from a table.

## Data Records

AncientMetagenomeDir (https://github.com/SPAAM-workshop/AncientMetagenomeDir) and AncientMetagenomeDirCheck (https://github.com/SPAAM-workshop/AncientMetagenomeDirCheck) are both maintained on GitHub. AncientMetagenomeDir has regular periodic releases, each of which has a release-specific DOI assigned via the Zenodo long-term data repository. Both the collection and tools are archived in the Zenodo repository with generalised DOIs: https://doi.org/10.5281/zenodo.3980833 and https://doi.org/10.5281/zenodo.4003826 respectively. The full workflow can be seen in Figure 3.

**Figure 3.**
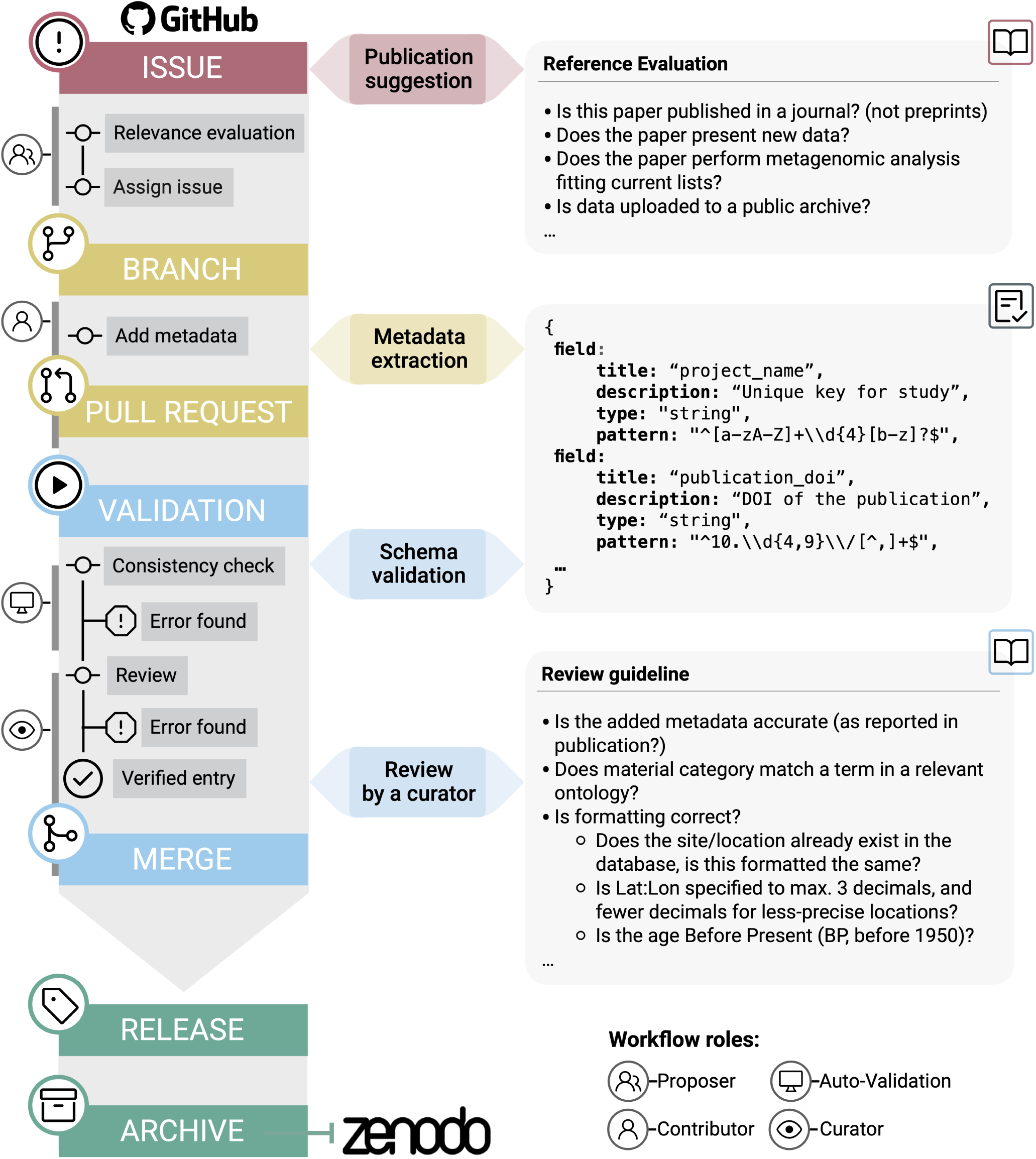
AncientMetagenomeDir submission and update workflow. The submission workflow is carried out on GitHub, and final releases archived at Zenodo. Submissions go through both automated computational validation and also human peer-review for consistency and accuracy.

## Technical Validation

All data entries to AncientMetagenomeDir undergo automated continuous-integration validation prior to submission into the protected main branch. These tests must pass before being additionally peer-reviewed by other member(s) of the community. Validation tests consist of regex patterns to control formatting of specified fields (e.g. DOIs, project IDs, date formats), and cross-checking of entries against controlled vocabularies defined in centralised JSON-format enum lists. Entries must also have valid sample accession IDs corresponding to shotgun metagenomic or genome-enriched sequence data uploaded to established and stable public archives.

## Usage Notes

Usage of the resource typically consists of copying or downloading the TSV file of interest for subsequent analysis using software such as Microsoft Excel, LibreOffice Calc, or R. The data table can be subsequently sorted or queried to identify datasets of interest. It should be noted that certain metadata fields (e.g., sample_age, latitude, and longitude) are approximate and do not provide *exact* values; rather, if exact values for these fields are required, they must be retrieved from the original publication. All selected data retrieved using AncientMetagenomeDir and used in subsequent studies should be cited using the original publication citation as well as AncientMetagenomeDir.

Retrieval of sequencing data using sample accession codes can be achieved manually via a given archive’s website, or via archive-supplied tools (e.g., Entrez Programming Utilties for NCBI’s SRA (https://github.com/enasequence/enaBrowserTools), or enaBrowserTools for EBI ENA (https://github.com/enasequence/enaBrowserTools).

Contributions to the tables are also facilitated by extensive step-by-step documentation on how to use GitHub and AncientMetagenomeDir, the locations of which are listed on the main README of the repository and the associated wiki page.

## Code availability

R notebooks used for generating images can be found at 10.5281/zenodo.4011751. Code for validation of the dataset (with version 1 used for the first release of AncientMetagenomeDir) can be found at https://github.com/SPAAM-workshop/AncientMetagenomeDirChec and https://doi.org/10.5281/zenodo.4003826.

## Acknowledgements

We would like to thank the wider SPAAM community (spaam-workshop.github.io) for their input in developing the project. J.A.F.Y., A.A.V., I.V., M.B. A.H. and C.W. acknowledge the Max Planck Society for financial support. J.A.F.Y. is partly supported by grant ERC-2015-StG 678901-FoodTransforms (to Philipp W. Stockhammer, Ludwig Maximilian University, Germany). B.C. is supported by grant ERC-2014-ADG 670518 (to V. Gaffney, University of Bradford, United Kingdom). A.J.V. is supported by Carlsbergfondet Semper Ardens grant CF18-1109 (to M. Thomas P. Gilbert, University of Copenhagen, Denmark). A.H. is partly supported by the Deutsche Forschungsgemeinschaft (DFG, German Research Foundation) under Germany’s Excellence Strategy—EXC 2051—Project-ID 390713860 (to C. Warinner, Friedrich Schiller University, Germany). E.J.G is supported by Arts & Humanities Research Council (grant number AH/N005015/1) and Natural History Museum (London, United Kingdom). M.J.B.-L. is supported by grant Wellcome Trust Seed Award in Science 208934/Z/17/Z, and by project IA201219 PAPIIT-DGAPA-UNAM (to María Ávila Arcos, LIIGH, Mexico). M.S. is supported by grant ERC-CoG 771234 PALEoRIDER (to Wolfgang Haak, Max-Planck-Institute for the Science of Human Hisory, Germany). A.S.G is supported by NSF GRFP Grant No. DGE1255832 (any opinions, findings, and conclusions or recommendations expressed in this material are those of the author(s) and do not necessarily reflect the views of the National Science Foundation). S.L.R. is supported by NIH Genetics and Regulation Training Grant 5T32GM007197-46. J.H. is supported by the Social Sciences and Humanities Research Council (Canada). I.V., M.B., and C.W. are supported by Werner Siemens Stiftung (Paleobiochemistry) (to C. Warinner, Leibnitz Institute for Natural Product Research and Infection Biology, Germany).

## Author contributions statement

J.A.F.Y and C.W. conceptualised the project. J.A.F.Y designed the project and infrastructure with input from all co-authors. M.B. developed software. J.A.F.Y., A.A.V., I.M.V., B.C., A.J.V., M.J.B-L., A.F.-G., E.J.G., S.L.R., P.D.H., M.A.S., A.H., A.S.G., J.H., A.F.A., and C.W. acquired data. J.A.F.Y drafted the manuscript with input from all co-authors.

